# Transient interdomain interactions shape the conformational ensemble governing RNA recognition by the tandem RRMs of Sex-lethal

**DOI:** 10.64898/2026.08.06.743427

**Authors:** Julia Meyer, Kristian Schweimer, Paul Matzner, Shohei Yoshida, Andrea Lomoschitz, Sandra Augsten, Bernd Simon, Po-Chia Chen, Janosch Hennig

## Abstract

RNA recognition motif (RRM) proteins frequently contain multiple RNA-binding domains connected by flexible linkers, yet the contribution of transient interdomain interactions to RNA recognition remains incompletely understood. Here, we investigated the structural organization of the tandem RRMs of the *Drosophila melanogaster* splicing regulator Sex-lethal (Sxl) using solution NMR spectroscopy in combination with rational protein engineering, restrained docking and RNA-binding studies. Progressive extension of the native interdomain linker resulted in a gradual decrease in rotational coupling between the two RRMs and continuous chemical shift changes, demonstrating that the RNA-free protein samples a dynamic conformational ensemble rather than behaving as two independently tumbling domains. NMR-guided docking identified a compact arrangement compatible with the experimental data and suggested a transient interface partially overlapping the RNA-binding surfaces. Surprisingly, a mutant designed to weaken this interface produced the opposite effect: instead of increasing interdomain mobility, it exhibited enhanced rotational coupling while remaining natively folded, indicating a redistribution of the conformational ensemble rather than disruption of the domain architecture. Both linker extension and the mutant reduced RNA-binding affinity, and the mutant additionally diminished sequence discrimination, demonstrating that perturbations shifting the conformational equilibrium in either direction compromise RNA recognition. Together, our results demonstrate that RNA recognition by Sxl is governed not by a single apo structure but by a finely balanced conformational ensemble, and that perturbing this equilibrium in either direction compromises high-affinity and sequence-selective RNA binding.

## Introduction

The Sex-lethal (Sxl) protein of *Drosophila melanogaster* is a female-specific RNA-binding protein (RBP) that plays a central role in regulating sex determination and dosage compensation [1]. As a master regulator, Sxl controls alternative splicing of its own pre-mRNA and several additional transcripts, most notably transformer (tra) pre-mRNA, thereby directing female-specific developmental programs [2]. Beyond its role in splicing, Sxl also acts as a potent translational repressor. Its best-characterized translational target is male-specific lethal 2 (msl2) mRNA, which encodes the limiting component of the male-specific lethal (MSL) dosage compensation complex [3, 4]. In male flies, the MSL complex mediates a two-fold upregulation of transcription from the single X chromosome to compensate for chromosome dosage. In females, inappropriate expression of msl2 is lethal, making translational repression by Sxl essential for viability.

Translational repression of msl2 is mediated through the cooperative assembly of a multi-protein ribonucleoprotein complex. Sxl and the RBP Upstream-of-N-Ras (Unr) bind cooperatively to conserved poly(U) stretches located in both the 3′ untranslated region (3′UTR) and the 5′ untranslated region (5′UTR) of msl2 mRNA (Figure 1A) [5–8]. This dual-site recognition establishes a robust repression mechanism in which 3′UTR-bound Sxl/Unr complexes prevent ribosome recruitment, while additional Sxl molecules bound to the 5′UTR inhibit translation initiation by ribosomes that escape the 3′UTR-dependent control. Efficient repression further requires the RBP Hrp48, which binds adjacent to the Sxl/Unr complex and stabilizes assembly of the quaternary messenger ribonucleoprotein (mRNP) complex [9, 10]. Structural and biophysical studies combining X-ray crystallography, NMR spectroscopy, small-angle scattering and single-molecule fluorescence microscopy have provided detailed insights into the architecture, assembly pathway and cooperative RNA recognition of the Sxl/Unr/Hrp48 complex, revealing both canonical and non-canonical protein-RNA interactions that underlie its remarkable sequence specificity [10–16]. Nevertheless, despite extensive characterization of the assembled complex, comparatively little is known about the structural organization of RNA-free Sxl itself and how this organization contributes to RNA recognition.

**Figure 1:**
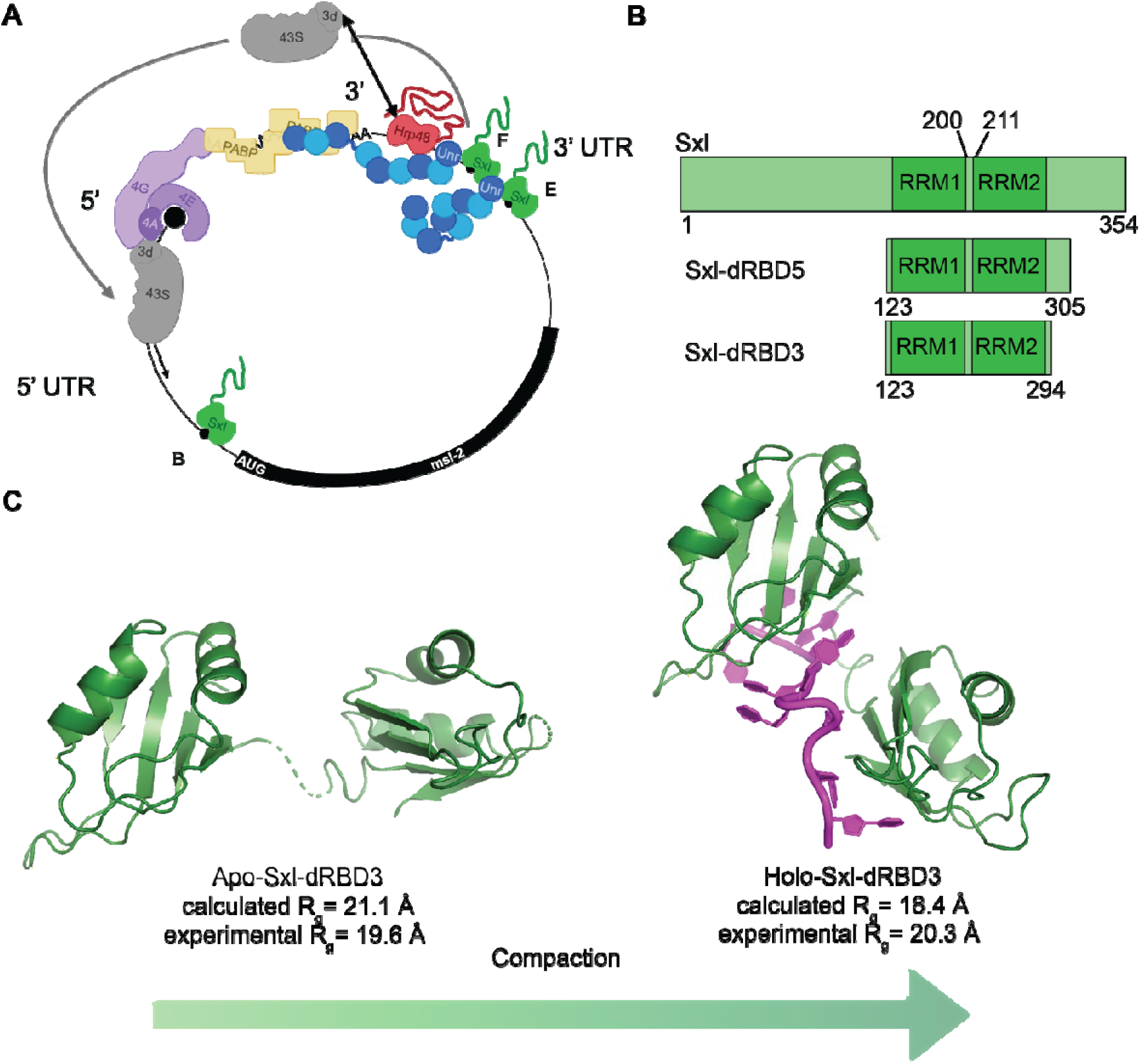
A: Schematic of Sxl-mediated translation repression of msl2-mRNA. B: Domain arrangement of Sxl. Sxl-dRBD3 is used in this study. C: The radius of gyration (R_g_) calculated from available crystal structures (apo-Sxl PDB 3SXL, Holo-Sxl PDB 1B7F) should decrease. The experimentally determined R_g_ shows that apo-Sxl is more compact than expected from the crystal structure.

Sxl contains two highly conserved RNA recognition motifs (RRM1 and RRM2) connected by a short, ten-residue linker (Figure 1B). The intrinsically disordered regions flanking the tandem RRMs are largely dispensable for RNA binding and translational repression, except for residues C-terminal to RRM2 (294–311), which adopt a non-canonical α-helix upon RNA binding [10]. Consequently, the Sxl-dRBD3 construct (residues 123–294), comprising only the tandem RRMs, faithfully recapitulates RNA binding and formation of the Sxl/Unr complex and therefore provides an excellent model system for investigating the structural basis of RNA recognition.

Tandem RRM domains represent one of the most common architectures found in RNA-binding proteins and frequently enhance RNA recognition through cooperative binding, modulation of affinity and increased sequence specificity [17–20]. In many multidomain RBPs, the relative orientation of individual RRMs is not fixed but instead samples multiple conformational states that are shifted upon RNA binding. In particular, work by Sattler et al. studied the splicing factor U2AF2 and demonstrated that transient interdomain interactions and linker-mediated conformational selection modulate recognition of strong and weak polypyrimidine tracts, thereby coupling domain organization directly to RNA-binding affinity and specificity [21–23]. These observations suggest that transient interdomain interactions may represent a general mechanism by which tandem RRMs regulate RNA recognition, although the structural principles governing this process are likely to differ between individual proteins.

Consistent with this idea, available structural information suggests that the relative arrangement of the two Sxl RRMs changes during RNA binding. The crystal structure of apo-Sxl (PDB: 3SXL) [24] adopts an extended conformation in which the two RRMs are spatially separated and do not form detectable interdomain contacts, whereas RNA-bound Sxl (PDB: 1B7F) [25] exhibits a substantially more compact arrangement in which both RRMs cooperatively engage the RNA substrate (Figure 1C). However, our previous small-angle X-ray scattering (SAXS) analysis demonstrated that the radius of gyration of RNA-free Sxl in solution is significantly smaller than predicted from the apo crystal structure, suggesting that the protein samples more compact conformations already in the absence of RNA [26]. Whether these conformations arise from transient interdomain interactions and whether such interactions contribute functionally to RNA recognition remained unknown.

Here, we combine solution NMR spectroscopy, rational protein engineering, restrained docking and quantitative RNA-binding measurements to investigate the conformational organization of the tandem RRMs of Sxl in solution. By systematically perturbing the relative arrangement of the two domains through linker extension and NMR-guided mutagenesis, we examine how changes in interdomain dynamics influence RNA binding. Our results demonstrate that RNA-free Sxl populates a dynamic conformational ensemble maintained by transient interdomain interactions and show that perturbing this equilibrium in either direction compromises high-affinity and sequence-selective RNA recognition.

## Results

### Linker extension progressively increases the independent motion of the two Sxl RRM domains

To determine whether the two RRM domains of Sxl interact in solution, we examined how extension of the interdomain linker affects their rotational dynamics. For two domains connected by a flexible linker but lacking appreciable interdomain contacts, increasing linker length is expected to promote progressively more independent domain motion. This relationship was previously characterized using artificial tandem GB1 constructs connected by linkers of different lengths. In that system, the rotational correlation times, τ*_c_*, of the individual domains decreased strongly upon initial linker extension and approached a plateau at longer linker lengths, consistent with largely independent domain tumbling [27].

Previous ^15^N spin relaxation measurements of wild-type Sxl-dRBD3, hereafter Sxl-WT, had indicated that its two RRM domains do not tumble independently. Sxl-dRBD3 has a molecular mass of 19.6 kDa, with each RRM accounting for approximately 9.5 kDa. At a protein concentration of 250 µM, Sxl-WT displayed an apparent rotational correlation time of 10.9 ns[28]. By comparison, the isolated RRM1 and RRM2 domains have a τ*_c_* of approximately 5.5 ns, as expected for a globular domain of about 10 kDa. The elevated value observed for the tandem construct could therefore reflect coupling of the rotational motion of the two RRMs. However, an intermolecular contribution could not be excluded, particularly because the apparent τ*_c_* increased to 15.1 ns at an Sxl-WT concentration of 1 mM [28].

To perturb potential intramolecular contacts without altering the folded RRM domains, we extended the native linker by inserting glycine-serine-rich sequences between G204 and G205. This generated three constructs containing 10, 20, or 25 additional residues, termed Sxl-GS10, Sxl-GS20, and Sxl-GS25, respectively (Figure 2A).

**Figure 2:**
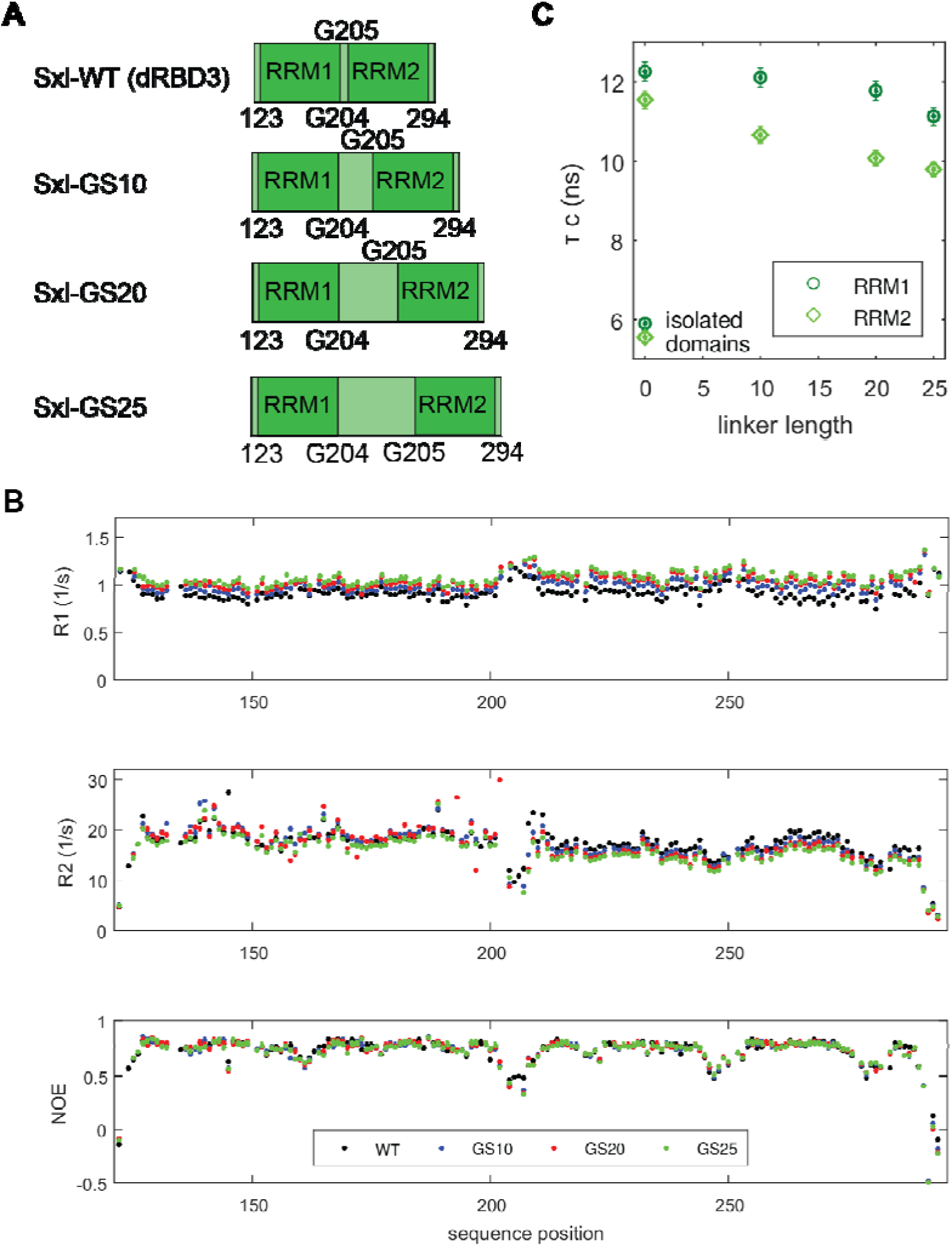
Linker extension progressively reduces interdomain coupling between the two RRMs of Sxl. A: Schematic representation of the Sxl-dRBD3 wild-type construct (Sxl-WT) and the glycine-serine linker extension variants Sxl-GS10, Sxl-GS20 and Sxl-GS25. The native ten-residue linker connecting RRM1 and RRM2 was progressively extended by insertion of glycine-serine repeats (see methods). B: Backbone ^15^N spin relaxation parameters recorded at 700 MHz for Sxl-WT (black), Sxl-GS10 (blue), Sxl-GS20 (red) and Sxl-GS25 (green). Residue-specific longitudinal relaxation rates (R_1_), transverse relaxation rates (R_2_) and heteronuclear ^1^H-^15^N NOEs are plotted as a function of sequence position. Overall, the relaxation profiles remain highly similar among all constructs, indicating that linker extension does not perturb the folded structures of the individual RRMs. The GS-linker residues are not assigned and therefore excluded from the figure and analysis. C: Rotational correlation times (τ_c_) determined separately for RRM1 and RRM2 as a function of linker length. The isolated RRM1 and RRM2 domains are shown for comparison at a linker length of zero. Progressive linker extension results in a gradual decrease in τ_c_ for both domains, consistent with reduced interdomain coupling. However, even the GS25 construct remains markedly distinct from the isolated domains, indicating that the two RRMs do not behave as fully independent tumbling units and supporting the presence of residual transient interdomain interactions.

We recorded ^15^N longitudinal and transverse spin relaxation rates at multiple magnetic-field strengths and used these data to estimate the rotational correlation times of the individual domains with ROTDIF [29] (Figure 2B,C and Supplementary Figure 1).

The rotational correlation times decreased progressively with increasing linker length (Figure 2C). This behavior is consistent with increasing motional independence of the two RRMs as the linker is extended. However, unlike the non-interacting tandem GB1 reference system, the Sxl constructs did not reach a clear plateau, even after insertion of 25 additional residues. Moreover, the values remained above that expected for independently tumbling RRM domains. Thus, linker extension weakens the coupling between the two domains but does not fully reproduce the behavior of isolated RRMs.

Together with the previously reported SAXS data, these results support an ensemble in which RRM1 and RRM2 transiently associate in RNA-free Sxl-WT. Increasing linker length appears to shift this ensemble toward conformations with greater motional independence rather than producing an abrupt transition between fully associated and fully independent domains.

### A ubiquitin-RRM2 control reproduces the behavior expected for non-interacting domains

To establish whether the linker-dependent changes observed for Sxl were specific to interactions between RRM1 and RRM2, we generated a control construct in which RRM1 was replaced by ubiquitin (Ub-RRM2). Ubiquitin is a compact globular domain of comparable, although slightly lower, molecular mass and was not expected to form a specific interface with Sxl RRM2. The hybrid constructs therefore retained a two-domain architecture while eliminating the native RRM1–RRM2 surface.

Four Ub-RRM2 constructs were produced with linker insertions of 3, 5, 10, or 15 glycine– serine residues (Figure 3A). Because ubiquitin and RRM2 differ in size, independently tumbling domains were expected to display distinct rotational correlation times. Furthermore, linker extension should produce a rapid decrease in the apparent rotational coupling followed by a plateau once the domains moved largely independently.

**Figure 3:**
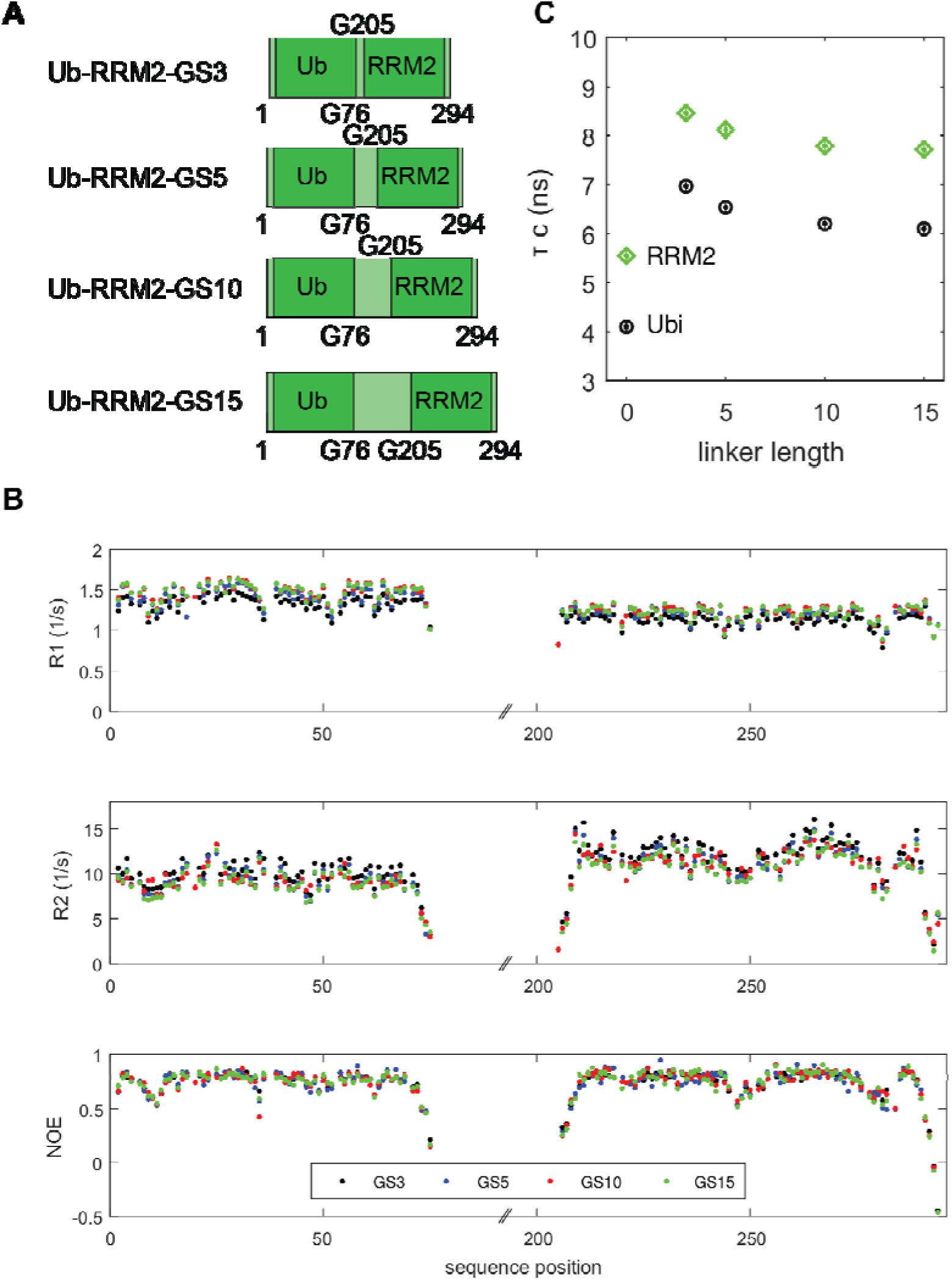
Ubiquitin–RRM2 fusion proteins exhibit progressively reduced rotational coupling upon linker extension. A: Schematic representation of the ubiquitin–RRM2 (Ub-RRM2) control constructs. Ubiquitin (Ub; residues 1–76) was fused to Sxl-RRM2 (residues 205–294) through glycine-serine linkers of increasing length, generating the constructs Ub-RRM2-GS3, Ub-RRM2-GS5, Ub-RRM2-GS10 and Ub-RRM2-GS15. B: Backbone ^15^N spin relaxation parameters recorded at 700 MHz for Ub-RRM2-GS3 (black), Ub-RRM2-GS5 (blue), Ub-RRM2-GS10 (red) and Ub-RRM2-GS15 (green). Residue-specific longitudinal relaxation rates (R_1_), transverse relaxation rates (R_2_) and heteronuclear ^1^H-^15^N NOEs are plotted as a function of sequence position. The relaxation profiles are highly similar among all constructs, indicating that linker extension does not perturb the folded structures of either ubiquitin or RRM2. The GS-linker residues are not assigned and therefore not shown. C: Rotational correlation times (τ_c_) determined separately for ubiquitin and RRM2 as a function of linker length. Both domains exhibit progressively decreasing τ_c_ values with increasing linker length, indicating that rotational coupling was mediated by a short linker and not interdomain interactions. In contrast to the Sxl tandem RRMs (Figure 2), relatively short linker extensions are sufficient to substantially decrease the apparent hydrodynamic coupling of the domains, consistent with independent tumbling behaviour of the Ub–RRM2 fusion proteins. The more pronounced decrease in τ_c_ compared to the Sxl linker variants supports the conclusion that the persistent coupling observed for the Sxl tandem RRMs cannot be explained by linker length alone. The τ_c_ values for isolated RRM2 and ubiquitin [30] domains are shown in the left for reference.

Consistent with these expectations, ^15^N spin relaxation measurements yielded different rotational correlation times for ubiquitin and RRM2 within the same construct (Figure 3B,C and Supplementary Figure 2). The linker-length dependence also closely resembled that previously observed for the non-interacting tandem GB1 constructs: the correlation times decreased rapidly upon linker extension and subsequently approached a plateau 40~50% above the isolated domain values. This is comparable to the 30% observed by Walsh et al. [27] and thus indicate that ubiquitin and RRM2 undergo largely independent rotational motion and validate the use of linker-length-dependent relaxation as a means of distinguishing interacting from non-interacting domain pairs.

The contrasting behavior of the ubiquitin-RRM2 controls and the Sxl linker variants therefore support the presence of transient intramolecular contacts between the native Sxl RRM domains. In the native tandem construct, these contacts maintain partial motional coupling even when the linker is substantially extended.

### Linker-dependent chemical shift changes identify a putative RRM1-RRM2 interface

We next sought to identify residues contributing to the interdomain interaction. To this end, we compared ^1^H,^15^N-HSQC spectra of Sxl-WT, the three linker-extension variants, and the isolated RRM1 and RRM2 domains (Figure 4A and Supplementary Figure 3A). Chemical shift differences relative to Sxl-WT were quantified in a residue-specific manner and plotted analogously to a chemical shift perturbation analysis (Figure 4B and Supplementary Figure 3B).

**Figure 4:**
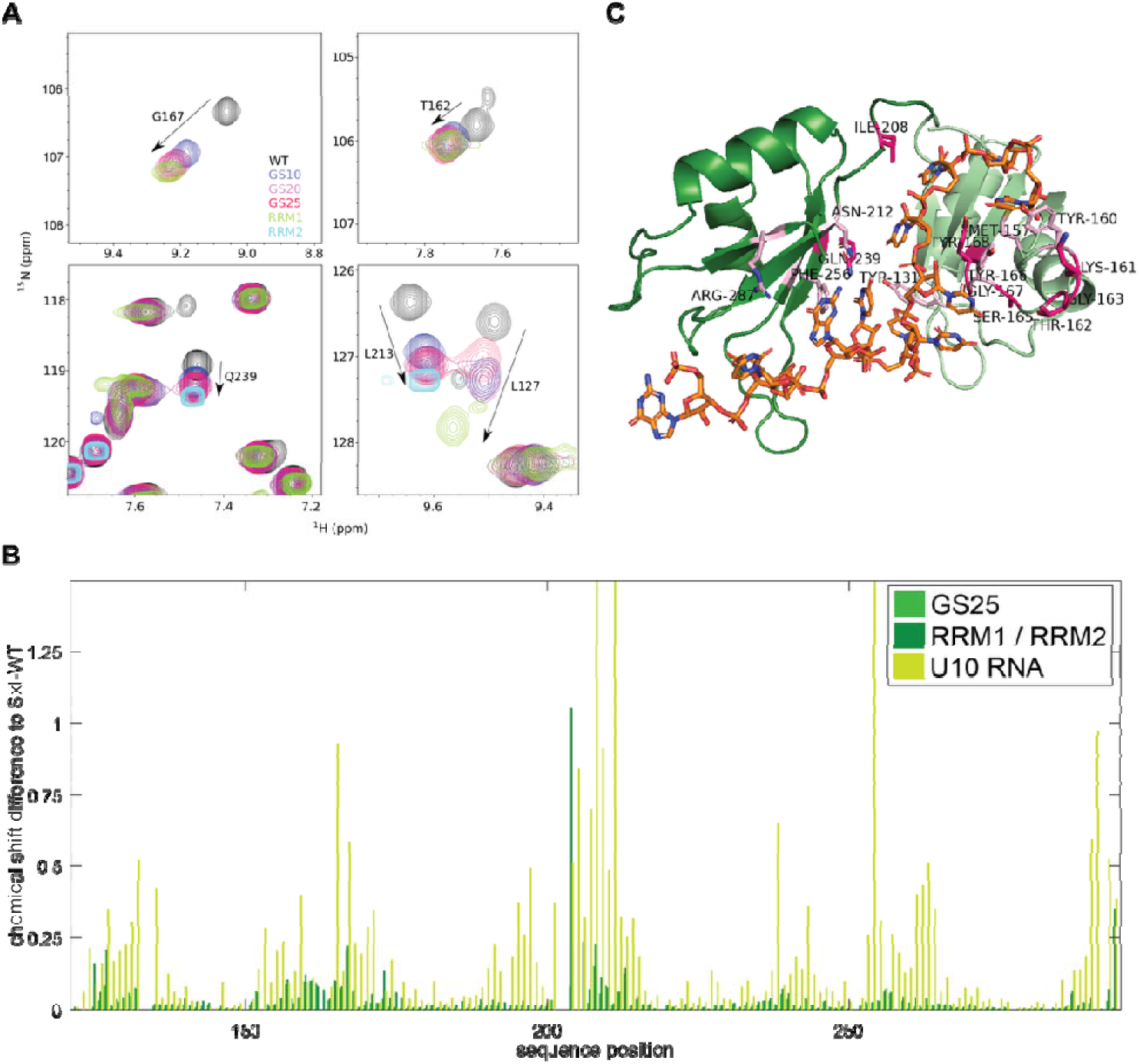
Linker extension induces chemical shift changes that map to a putative interdomain interface overlapping the RNA-binding surface. A: Representative regions of ^1^H-^15^N HSQC spectra illustrating linker length-dependent chemical shift changes. Spectra of Sxl-WT, Sxl-GS10, Sxl-GS20, Sxl-GS25, and the isolated RRM1 and RRM2 domains are overlaid as indicated. Selected resonances exhibit progressive movement from their positions in Sxl-WT toward those observed for the isolated domains as linker length increases, consistent with a gradual redistribution of conformational populations upon weakening interdomain coupling. B: Combined amide chemical shift perturbations (CSPs) plotted as a function of sequence position. CSPs were calculated for Sxl-GS25 relative to Sxl-WT (green), the isolated RRM domains relative to Sxl-WT (dark green), and RNA-bound Sxl-WT relative to free Sxl-WT (light green). Residues exhibiting the largest perturbations cluster on distinct surfaces of both RRMs and partially overlap with residues affected by RNA binding. C: Mapping of residues exhibiting significant CSPs in the GS25 versus WT comparison onto the crystal structure of Sxl bound to a U10-RNA (PDB: 1B7F). RRM1 and RRM2 are shown in light and dark green, respectively, and the RNA is shown as sticks. Residues displaying significant CSPs upon linker extension are highlighted and labelled. Residues that are also perturbed upon RNA binding are shown in light pink, whereas residues affected by linker extension but not by RNA binding are shown in magenta. The spatial clustering of these residues identifies a putative interdomain interaction surface that partially overlaps with the RNA-binding interface.

A subset of resonances shifted progressively as the linker was extended. For these residues, the peak positions followed a trajectory from Sxl-WT, through Sxl-GS10 and Sxl-GS20, toward Sxl-GS25 and the corresponding isolated RRM domains. The shifts were therefore not random consequences of inserting a flexible sequence into the linker. Instead, they are consistent with a gradual change in the population of conformations contributing to the observed spectrum.

Even for Sxl-GS25, however, the affected resonances did not fully coincide with those of the isolated domains. This observation agrees with the relaxation data and suggests that the longest linker variant retains residual interdomain coupling or populates an ensemble that remains distinct from the completely isolated RRMs.

Residues exceeding the significance threshold (average plus standard deviation) were located in both domains. In RRM1, these include Y131, M157, Y160, K161, T162, G163, S165, Y166, G167, and Y168. In RRM2, significant perturbations were observed for I208, N212, Q239, F256, R287, and E291 (Figure 4C). These residues define candidate regions involved either directly in the interdomain interface or indirectly through conformational changes associated with linker extension.

Notably, several of the affected residues, including Y131, M157, Y160, Y166, Y168, N212, F256, and R287, also contact RNA in available Sxl-RNA structures (PDBs: 1B7F, 4QQB) [12, 25]. The putative interdomain surface therefore overlaps substantially with the canonical RNA-binding surfaces of the RRMs. This overlap suggests that interdomain association and RNA recognition are structurally coupled, although chemical shift perturbations alone cannot distinguish direct interface contacts from allosteric or ensemble-dependent effects.

### NMR-guided docking supports a compact RNA-free arrangement of the tandem RRMs

Having identified residues affected by linker extension, we asked whether they could define a physically plausible interface between RRM1 and RRM2. We therefore performed HADDOCK calculations using the individual RRM structures as separate coordinate files derived from the RNA-bound Sxl structure and the significant chemical shift perturbations as ambiguous interaction restraints (see Methods) [31, 32].

The docking calculation produced five clusters when structures were grouped using a fraction-of-common-contacts cutoff of 0.6 and a minimum cluster size of four structures (Figure 5A). Cluster 1 was the largest, yielded significantly more favorable interface energies than other clusters, and included the lowest energy structure among the 200 water refined models (Figure 5A,B). In this cluster, the C-terminus of RRM1 and the N-terminus of RRM2 were positioned sufficiently close to permit connection by the unresolved native linker following modest rearrangement of its flexible residues.

**Figure 5:**
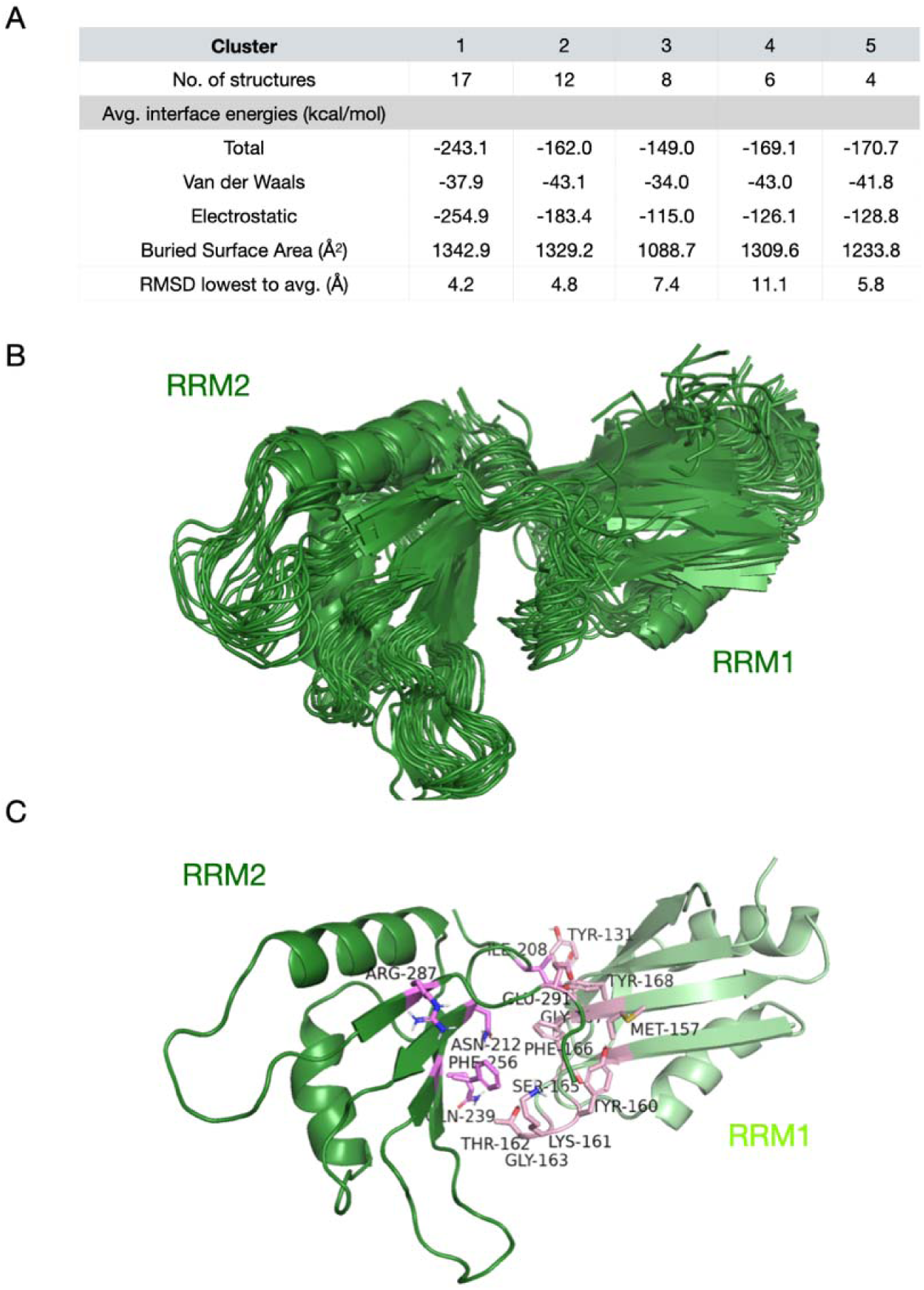
HADDOCK-based structure modelling of a potential RRM1-RRM2 interface based on chemical shift perturbation data (Sxl-WT vs. Sxl-GS25). A: Statistics of the HADDOCK run. RMSD lowest to average is a comparison of the backbone root mean square deviation between the lowest energy structure of each cluster with the overall average structure. B: Sxl-WT structure ensemble of the 17 structures of the best cluster (cluster 1) of the HADDOCK run. C: Lowest energy structure of cluster 1. Highlighted are active interaction restraints, which drove the HADDOCK run identified by NMR chemical shift perturbations between Sxl-WT and Sxl-GS25).

The models place RRM2 against the β-sheet surface of RRM1, thereby partially occluding the canonical RNA-binding surface (Figure 5C). Several residues used as active docking restraints are positioned at the resulting interface. This arrangement provides a structurally plausible explanation for the linker-dependent relaxation and chemical shift perturbation data and is consistent with an RNA-free ensemble containing compact conformations.

The HADDOCK ensemble should not, however, be interpreted as a uniquely determined structure of apo-Sxl-WT. Chemical-shift perturbations provide limited spatial information and can arise from both direct contacts and indirect conformational effects. The model therefore represents one experimentally compatible arrangement that rationalizes the observed perturbations and guides subsequent mutational analysis.

### Design of a mutant intended to perturb the proposed interdomain interface

We next attempted to perturb the proposed interface while minimizing direct effects on RNA recognition. This was challenging because the residues affected by linker extension overlap extensively with the RNA-binding surfaces of both RRMs. Complete separation of interdomain and RNA-binding determinants was therefore unlikely to be achievable by straightforward point mutagenesis.

Based on the chemical shift analysis and the HADDOCK model, we selected T162, Y168, R258, and E291 as candidate residues. T162 and E291 displayed significant linker-dependent perturbations, whereas R258 was positioned near the center of the modeled interface despite not exhibiting a comparably large chemical shift change. Y168 contributes with its hydroxyl group a hydrogen bond to RNA in the bound structure but does not form the extensive aromatic stacking interaction characteristic of several other RNA-contacting residues. We therefore reasoned that its mutation might be less disruptive to RNA binding than mutation of the principal stacking residues.

The four substitutions T162G, Y168A, R258A, and E291G were combined in a single construct, termed the TYRE mutant (Figure 6A). The mutant was designed to weaken the proposed RRM1–RRM2 interface and was initially expected to display increased motional independence of the two domains.

**Figure 6:**
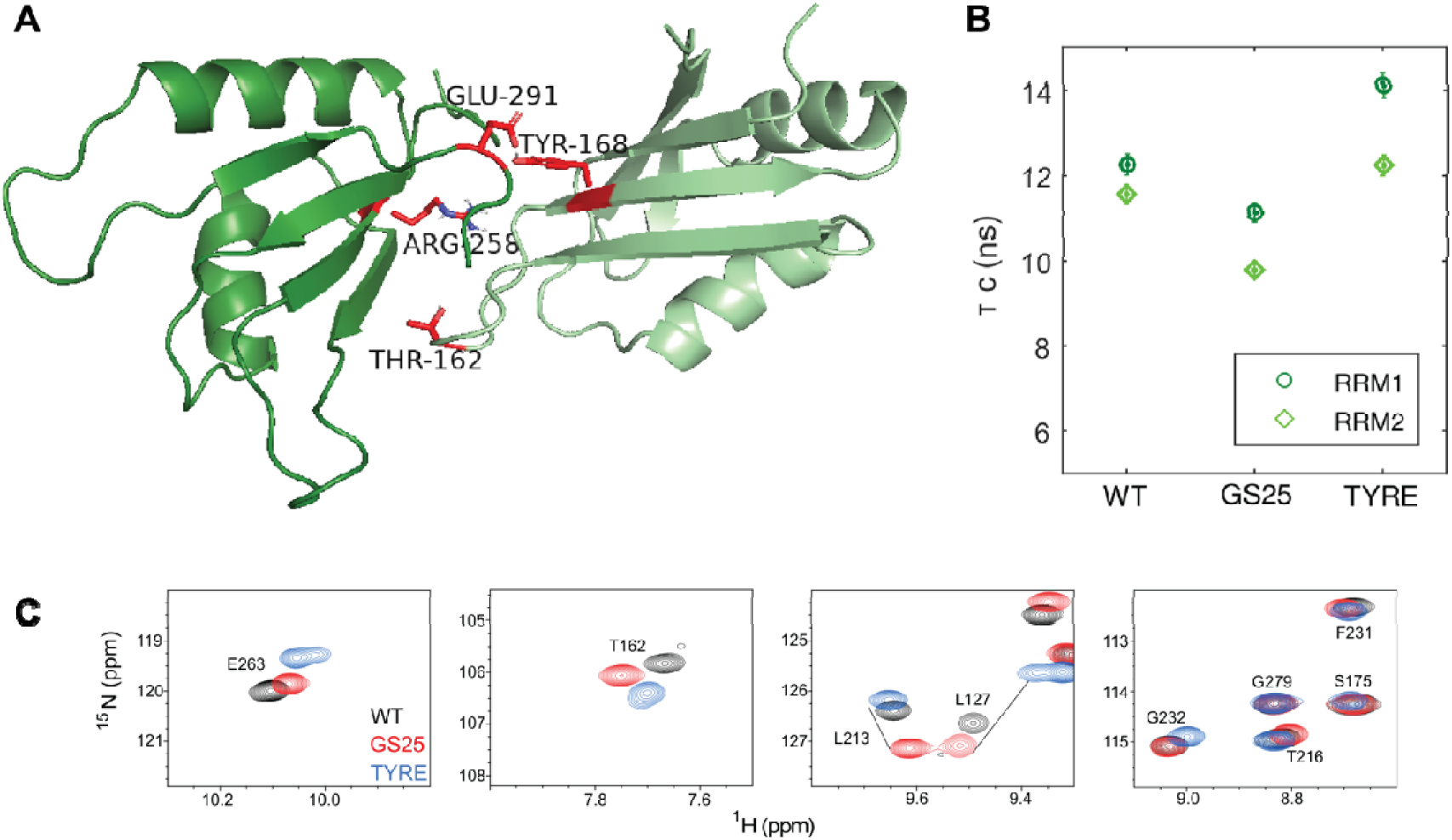
The TYRE mutant reshapes the conformational ensemble of the Sxl tandem RRMs. A: Lowest-energy structure of the highest-ranking HADDOCK cluster generated using residues identified by chemical shift perturbation analysis as ambiguous interaction restraints. RRM1 and RRM2 are shown in light and dark green, respectively. The four residues selected for mutational analysis (TYRE) are highlighted in red. B: Rotational correlation times (τ_c_) determined separately for RRM1 and RRM2 in Sxl-WT, Sxl-GS25 and the TYRE mutant. As shown in Figure 2, linker extension in GS25 reduces rotational coupling between the two domains, resulting in lower τ_c_ values. In contrast, the TYRE mutant exhibits increased τ_c_ values relative to both GS25 and WT, indicating enhanced rotational coupling between the two RRMs despite the disruption of residues predicted to participate in the putative interdomain interface. C: Zoom-ins of ^1^H-^15^N HSQC spectra of Sxl-WT (black), Sxl-GS25 (red) and the TYRE mutant (blue). The spectra exhibit excellent chemical shift dispersion and comparable peak patterns, demonstrating that all three proteins are properly folded (Supplementary Figure 4B). While the TYRE mutant displays substantial chemical shift changes relative to WT and GS25, the overall spectral quality indicates that the observed effects arise from changes in the conformational ensemble rather than global unfolding or misfolding of either RRM.

### Mutational perturbation of the proposed interface produces a distinct conformational ensemble

To test this hypothesis, we first determined the rotational correlation times of the TYRE mutant using ^15^N spin relaxation experiments (Supplementary Figure 4A). Contrary to our expectation, the mutant did not display the reduced rotational coupling observed for the GS variants. Instead, both RRMs exhibited increased apparent rotational correlation times, with the largest effect observed for RRM1 (Figure 6B). Thus, rather than shifting the conformational ensemble toward increased domain independence, the TYRE mutant produced the opposite effect.

To further characterize the structural consequences of the mutations, we compared the ^1^H-^15^N HSQC spectrum of TYRE with those of Sxl-WT and Sxl-GS25. The spectrum of the mutant remained well dispersed, demonstrating that both RRMs retained their native folds (Supplementary Figure 4B). However, numerous resonances were shifted relative to both wild type and the linker variants, indicating that the mutations alter the conformational ensemble of the tandem RRMs rather than simply reproducing or extending the effect of linker extension (Figure 6C).

Interestingly, several resonances that shifted progressively from Sxl-WT toward Sxl-GS25 exhibited chemical-shift changes in the opposite direction in the TYRE mutant. Although these observations do not allow a direct structural interpretation, they are consistent with the notion that the mutations shift the conformational equilibrium toward a population distinct from both wild type and the linker extension variants. One possible explanation is that the compact state is stabilized or more highly populated in the TYRE mutant, resulting in increased rotational coupling between the two RRMs. While additional structural information will be required to distinguish between these possibilities, the relaxation and HSQC data clearly demonstrate that the TYRE mutations reshape the conformational landscape of Sxl-WT rather than simply disrupting a discrete interdomain interface.

### Reshaping the conformational ensemble compromises RNA recognition

Having established that the TYRE mutant alters the conformational landscape of the tandem RRMs, we next investigated how these changes affect RNA binding. We therefore compared RNA-induced chemical shift perturbations for Sxl-WT, the Sxl-GS25 linker variant, and the TYRE mutant using the high affinity U10-mer RNA ligand as well as a control RNA (U5C5) with reduced affinity (Figure 7A–C, Supplementary Figure 5).

**Figure 7:**
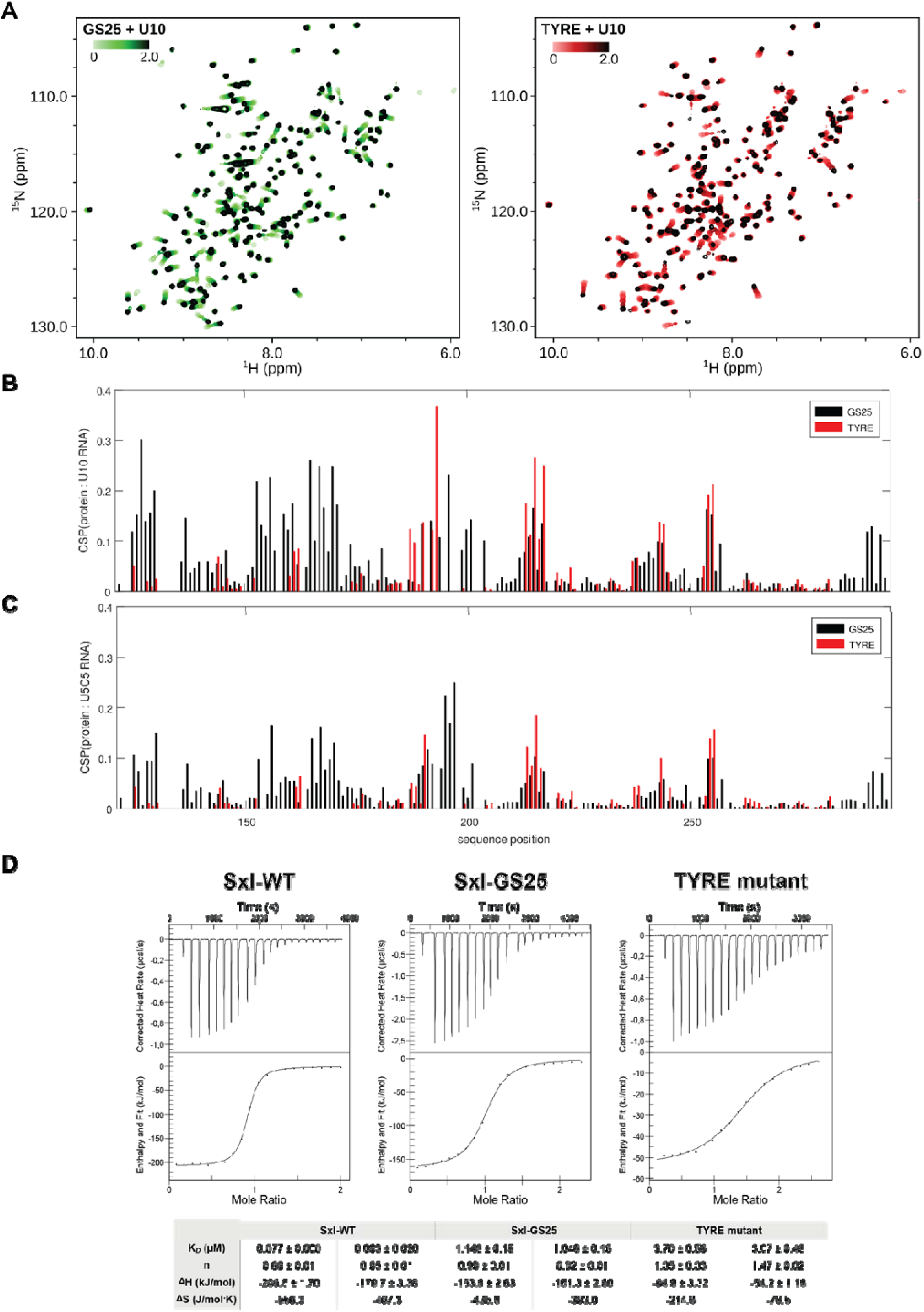
Perturbation of the conformational equilibrium compromises RNA recognition by the Sxl tandem RRMs. A: Overlay of 1H-15N HSQC spectra recorded during titration of Sxl-GS25 (left) and the TYRE mutant (right) with a U10 RNA oligonucleotide. Progressive chemical shift changes are observed for both proteins upon RNA addition, indicating RNA binding. The magnitude of the perturbations is generally reduced for the TYRE mutant compared to Sxl-GS25. B: Combined amide chemical shift perturbations (CSPs) induced by binding of U10 RNA plotted as a function of sequence position for Sxl-GS25 (black) and the TYRE mutant (red). Residues in both RRMs participate in RNA recognition; however, the overall magnitude of the CSPs is substantially reduced in the TYRE mutant, consistent with weaker RNA binding. C: Combined amide chemical shift perturbations induced by binding of a U5C5 RNA oligonucleotide plotted as a function of sequence position for Sxl-GS25 (black) and the TYRE mutant (red). Similar to the U10 titration, RNA-induced perturbations are markedly reduced in the TYRE mutant, indicating impaired recognition of both RNA substrates. In B and C, GS-linker residues of GS25 are not assigned and therefore not shown for easier comparison. D: Isothermal titration calorimetry (ITC) measurements of RNA binding by Sxl-WT, Sxl-GS25 and the TYRE mutant. Representative binding isotherms and corresponding fits are shown. Consistent with the NMR titration experiments, linker extension reduces RNA-binding affinity relative to the wild-type protein, whereas the TYRE mutant exhibits a further decrease in affinity. Together, these data demonstrate that perturbations shifting the conformational equilibrium in either direction compromise efficient RNA recognition by the tandem RRMs of Sxl.

Consistent with the relaxation data, the linker-extension mutant retained the characteristic RNA-binding mode of wild-type Sxl but exhibited generally smaller chemical shift perturbations, indicating reduced binding affinity while preserving sequence discrimination. In contrast, the TYRE mutant displayed substantially weaker perturbations throughout both RRMs together with a markedly altered perturbation pattern. The differences between the preferred U-rich RNA and the control RNA became considerably less pronounced, suggesting that the mutations not only reduces binding affinity but also sequence discrimination.

Interestingly, the reduction in RNA-induced chemical shift perturbations was most pronounced for RRM1, which also displayed the largest increase in apparent rotational correlation time in the relaxation experiments. Thus, the domain exhibiting the strongest increase in rotational coupling simultaneously contributes least to RNA binding. Although the present data do not establish a causal relationship, this correlation is consistent with the idea that increased interdomain coupling reduces the accessibility of the RNA-binding surface. However, the mutations in RRM1 could increase the transient RRM1-RRM1 intermolecular interactions observed earlier at high concentrations, also occluding the RNA binding residues [28].

To independently quantify these effects, RNA binding was further analyzed by isothermal titration calorimetry (ITC, Figure 7D). In agreement with the NMR titration experiments, progressive linker extension reduced RNA-binding affinity compared with Sxl-WT, whereas the TYRE mutant exhibited the weakest binding of all constructs. Thus, two independent perturbation strategies produced reciprocal changes in interdomain dynamics but converged on the same functional outcome: altering the native conformational ensemble compromises efficient RNA recognition.

Collectively, these findings demonstrate that the transient interdomain interaction is not merely a structural feature of apo-Sxl but contributes directly to the dynamic organization required for optimal RNA binding. Both increasing the conformational freedom of the two RRMs by linker extension and shifting the ensemble toward a more compact state by the TYRE mutations impair RNA recognition, indicating that efficient RNA binding depends on a finely balanced equilibrium between alternative interdomain conformations.

## Discussion

The tandem RNA recognition motifs (RRMs) of Sex-lethal have long served as a model system for understanding sequence-specific RNA recognition[18–20]. Structural studies of Sxl bound to RNA established the molecular basis for high-affinity recognition of U-rich sequences and demonstrated that both RRMs cooperate to achieve binding specificity [25]. In contrast, considerably less has been known about the structural organization of the RNA-free protein and whether interdomain interactions contribute to RNA recognition before ligand binding[24, 26]. In the present study, we combined linker engineering, solution NMR spectroscopy, restrained docking and mutational analysis to investigate the conformational organization of apo-Sxl. Together, our results demonstrate that the two RRMs do not behave as independent domains connected by a passive flexible linker. Instead, they populate a conformational ensemble in which transient interdomain interactions modulate the relative orientation and dynamics of the two domains and thereby influence RNA binding.

The progressive linker-extension strategy provides particularly compelling evidence for such transient interdomain interactions. Increasing linker length resulted in a gradual reduction of rotational coupling between the two RRMs, consistent with increasing motional independence of the domains. However, even the longest linker construct did not fully reproduce the behavior expected for independently tumbling domains, in clear contrast to the ubiquitin-RRM2 control and previously characterized artificial tandem proteins lacking detectable interdomain contacts [27]. Similarly, the accompanying chemical shift perturbations followed continuous trajectories rather than discrete transitions, indicating that linker extension shifts the populations of conformational states instead of simply disrupting a single defined interface. Together with our previous SAXS data [26], these observations support a model in which apo-Sxl samples a dynamic equilibrium between more compact and more extended conformations in solution.

The CSP-derived HADDOCK model provides a structurally plausible representation of one member of this ensemble. Importantly, we do not interpret this model as a unique structure of apo-Sxl. Chemical shift perturbations contain only limited spatial information and cannot distinguish direct intermolecular contacts from indirect conformational effects. Nevertheless, the resulting models place the RNA-binding surfaces of the two RRMs in close proximity, providing a straightforward explanation for the observed relaxation behavior and the linker-dependent chemical shift changes. Rather than defining a single structural solution, the docking calculations provide a structural framework that integrates the experimental observations and guided the subsequent mutational analysis.

Perhaps the most informative finding of this study emerged from the TYRE mutant. This construct was designed to weaken the proposed interdomain interface and was therefore expected to mimic the effects of linker extension. Instead, the opposite behavior was observed. The mutant exhibited increased rotational coupling between the two RRMs while maintaining well-dispersed HSQC spectra, demonstrating that both domains remain folded. Furthermore, several resonances shifted in a direction opposite to that observed upon linker extension, indicating that the TYRE mutations do not simply abolish the proposed interface but instead reshape the conformational ensemble sampled by the tandem RRMs. Although the present data do not allow us to define the resulting structural state, they are consistent with a redistribution of conformational populations, potentially favoring a more compact arrangement of the two domains. This reciprocal behavior of the linker variants and the TYRE mutant strongly argues that the observed relaxation changes reflect genuine alterations in interdomain organization rather than non-specific structural perturbations.

Remarkably, both perturbation strategies converged on the same functional outcome. Progressive linker extension reduced RNA-binding affinity, while the TYRE mutant, despite producing the opposite effect on interdomain dynamics, impaired RNA binding even more strongly. This observation suggests that efficient RNA recognition does not simply require weak or strong interdomain interactions. Instead, the wild-type protein appears to maintain a conformational equilibrium that is particularly well suited for RNA recognition. Perturbing this equilibrium in either direction, either by increasing the conformational freedom of the domains or by favoring a more tightly coupled arrangement, reduces RNA-binding affinity and, in the case of TYRE, also diminishes sequence discrimination. The data therefore suggest that transient interdomain interactions contribute to pre-organizing the tandem RRMs for efficient ligand recognition while retaining sufficient conformational flexibility to adopt the RNA-bound state.

This interpretation is consistent with an emerging view of multidomain RNA-binding proteins as dynamic systems whose functional properties arise from ensembles of interconverting conformations rather than single static structures. Several tandem RRM proteins, including Sxl, U2AF2, PTB, and HuR, undergo substantial rearrangements upon RNA binding, illustrating that relative domain orientation is frequently coupled to RNA recognition [20, 33–35]. Our work complements these studies by demonstrating how systematic perturbation of interdomain dynamics can be used to dissect the functional contribution of transient interdomain interactions in a well-defined model system.

Our work extends the above established concept by showing how systematic perturbation of interdomain dynamics can be used to dissect the functional contribution of these interactions in a well-defined model system. The reciprocal effects observed for linker extension and the TYRE mutant further illustrate that the energetic landscape governing domain organization is finely balanced and can be modulated in multiple ways, each producing distinct consequences for RNA recognition.

The importance of conformational ensembles for RNA recognition also highlights an important limitation of current machine learning-based protein structure prediction methods. Recent advances such as AlphaFold2 [36] and AlphaFold3 [37] have transformed structural biology by providing highly accurate models for many folded proteins and protein complexes. However, apart from mostly failing to accurately predict protein-RNA complexes [38], these approaches are designed to predict the most probable structural arrangement under a given set of constraints and generally do not describe the underlying conformational landscape sampled in solution. In particular, weakly populated states, transient interdomain interactions and rapidly interconverting domain arrangements, such as those observed for the tandem RRMs of Sxl, are largely invisible to current prediction methods. The present study therefore illustrates that experimentally characterizing conformational dynamics remains essential for understanding the molecular mechanisms of multidomain proteins. Rather than replacing experimental structural biology, predictive models and solution-state methods such as NMR spectroscopy provide complementary information, with the former defining plausible structural states and the latter revealing the dynamic equilibria that frequently determine biological function [39].

Our findings also illustrate the strengths of solution NMR spectroscopy for characterizing dynamic multidomain proteins. While static structural methods provide high-resolution snapshots of individual conformational states, relaxation measurements and titration experiments directly report on the dynamic equilibria that often underlie biomolecular function. In combination with rational protein engineering and complementary biophysical approaches such as ITC, these methods provide a powerful framework for connecting conformational dynamics with molecular function.

Several limitations of the present study should be acknowledged. First, the HADDOCK model represents one experimentally compatible arrangement within a dynamic conformational ensemble and should not be interpreted as a unique structural solution. Second, the residues selected for mutagenesis overlap partially with the RNA-binding surfaces of the RRMs, making it difficult to separate effects on interdomain interactions from direct effects on RNA recognition. Nevertheless, the reciprocal behavior of the linker variants and the TYRE mutant argues that changes in the conformational ensemble make a substantial contribution to the observed functional differences. Finally, while relaxation measurements clearly demonstrate altered interdomain dynamics, they do not directly resolve the individual conformational states or their exchange kinetics. Future studies employing complementary approaches such as residual dipolar couplings, paramagnetic relaxation enhancement measurements, or molecular dynamics simulations may further define the structural and energetic landscape of apo-Sxl.

In summary, our results demonstrate that the tandem RRMs of Sxl populate a dynamic conformational ensemble in solution in which transient interdomain interactions contribute directly to RNA recognition. Rather than behaving as two independently folded RNA-binding modules connected by a flexible linker, the domains cooperate through a finely balanced conformational equilibrium that optimizes RNA binding and sequence selectivity. We anticipate that similar principles will govern many multidomain RNA-binding proteins, highlighting the importance of considering conformational dynamics alongside static structural information when investigating mechanisms of RNA recognition.

## Methods

### Cloning

The DNA sequence encoding the tandem RNA recognition motifs of *Drosophila melanogaster* Sex-lethal (Sxl-dRBD3; residues 123-294) was part of the bacterial expression vector pET-M11as previously described[12]. Sxl linker extension variants were generated by insertion of glycine-serine repeats into the native interdomain linker using overlap-extension PCR. (GS10: Sxl-RRM1-G204-GGSGSGGGGS-G205-Sxl-RRM2, GS20: Sxl-RRM1-G204-GGSGGSGSGSGGSGSGGGGS-G205-Sxl-RRM2, GS25: Sxl-RRM1-G204-SGSGSGGSGGSGSGSGGSGSGGGGS-G205-Sxl-RRM2). The ubiquitin-RRM2 constructs were cloned in a similar way and comprised the following sequences: GS3: Ubiquitin-G78-GS-G205-Sxl-RRM2, GS5: Ubiquitin-G78-GGSGS-G205-Sxl-RRM2, GS10: Ubiquitin-G78-GGSGSGGGGS-G205-Sxl-RRM2, GS15: Ubiquitin-G78-SGSGSGGSGSGGGGS-G205-Sxl-RRM2. The TYRE mutant (T162G/Y168A/R258A/E291G) was generated by site-directed mutagenesis using a protocol previously described by Edelheit et al. [40] All resulting constructs encoded an N-terminal His_6_-tag followed by a TEV protease cleavage site and were verified by Sanger DNA sequencing.

### Protein expression and purification

All proteins were expressed as previously described [12]. In short, expression was induced in *Escherichia coli* Rosetta (DE3) cells. For isotopically labelled samples, cells were grown in M9 minimal medium supplemented with 0.5 g/L (^15^NH_4_)_2_SO_4_ and 2 g/L ^13^C-glucose as sole nitrogen and carbon sources, respectively if backbone assignment was needed or only (^15^NH_4_)_2_SO_4_. Cultures were grown at 37 °C to an OD600 of approximately 0.8 before protein expression was induced with 0.5 mM IPTG. Cells were incubated for an additional 16 h at 18 °C and harvested by centrifugation.

Cell pellets were resuspended in lysis buffer containing 50 mM sodium phosphate (pH 7.2), 500 mM NaCl, 1 mM DTT, and protease inhibitors. Cells were disrupted by French press, and the lysate was clarified by centrifugation at 10000 × g for 30 min. The soluble fraction was applied to a Ni^2+^-NTA affinity column (Cytiva), washed extensively with lysis buffer containing 25 mM imidazole, and eluted using 500 mM imidazole. The His_6_-tag was removed by overnight digestion with recombinant TEV protease during dialysis against 20 mM sodium phosphate (pH 7.2), 500 mM NaCl, and 1 mM DTT. Cleaved protein was separated from uncleaved material and TEV protease by a second Ni^2+^-NTA purification step. After size-exclusion chromatography on a S75 gel filtration column, the purified proteins were dialyzed against NMR buffer containing 10 mM sodium phosphate (pH 6.5), 50 mM NaCl, 1 mM DTT and subsequently concentrated to 400 μM using centrifugal concentrators (Millipore) and stored at −80 °C until use. For NMR experiments, 10% (v/v) D_2_O was added to the protein sample.

### NMR spectroscopy

All NMR experiments were performed at 298 K on Bruker Avance III or Avance III HD spectrometers operating at proton larmor frequencies of 500, 600, 700, 800, 900 and 1000 MHz equipped with cryogenically cooled triple-resonance probes (600, 700, 800, 900, 1000) or room temperature probes (500, 600, 700). Data were processed using NMRPipe [41] or in-house scripts and analyzed with NMRViewJ [42, 43] and in-house scripts.

Backbone ^15^N longitudinal (*R_1_*) and transverse (*R_2_*) relaxation rates were measured using standard Bruker pulse sequences. *R_1_*relaxation delays of 80(2x), 400, 800(2x), 1200(2x), 1600 and 2000(2x) ms (at 1000 MHz also 2400 and 2800 ms) and *R_2_* delays of 16, 32(2x), 48, 64, 80, 96, 112, 128, and 144 ms were recorded.

Peak intensities were extracted in NMRViewJ [42, 43] or PINT [44] and fitted to mono-exponential decay functions using nonlinear least-squares fitting. *R_1_, R_2_* rate errors were set to 3% in cases with lower values from data ftiing. Rotational correlation times (τ*_c_*) were estimated from spin relaxation data using ROTDIF [29]. Only data from residues located in regular secondary structure were used while excluding residues exhibiting significant internal dynamics or spectral overlap. Reported uncertainties represent standard deviations derived from the selected residue set.

### NMR chemical shift perturbation analysis

Two-dimensional ^1^H,^15^N-HSQC spectra were recorded for all protein variants under identical buffer conditions. Backbone resonance assignments were transferred from previously published assignments and verified by standard triple-resonance or ^15^N-edited NOESY experiments where required.

Combined amide chemical shift perturbations (CSPs) were expressed as weighted geometric average according to

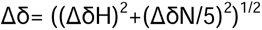

where ΔδH and ΔδN denote proton and nitrogen chemical shift differences, respectively [45]. Residues with broadened or overlapping resonances were excluded from the analysis. For RNA titrations, increasing amounts of RNA were added to uniformly ^15^N-labelled protein. Spectra were recorded after each addition and analyzed as described above.

### HADDOCK docking calculations

Protein-protein docking calculations were performed using the HADDOCK webserver [31]. Active residues were defined on the basis of residues exhibiting significant chemical shift perturbations upon linker extension, while neighbouring solvent-accessible residues were automatically assigned as passive residues. No explicit distance restraints derived from NOEs were included.

For each docking calculation, 1000 rigid-body structures were generated, followed by 200 structures in a semi-flexible simulated annealing and explicit solvent refinement step according to the standard HADDOCK protocol. The resulting models were clustered using an a fraction of common contacts (FCC) of 0.6 with a minimum cluster size of 4. Clusters were ranked according to the interface energies. The lowest energy structure in cluster 1 was additionally checked to confirm that the distance between RRM1 C-terminal and RRM2 N-terminal falls within the limits imposed by the 10-residue linker.

### Isothermal titration calorimetry

ITC experiments were performed using a AffinityITC with gold cell (TA instruments) at 25 °C. Protein and RNA samples were extensively dialyzed against identical buffer consisting of 10 mM sodium phosphate (pH 6.5), 50 mM NaCl and 1 mM DTT to minimize heats of dilution.

Typically, 16-49 μM RNA was placed in the sample cell and titrated with 129-493 μM protein using 20 injections of 2 μL each with a spacing of 150-300 s between injections and stirring rate of 125 rpm. The first injection was discarded during analysis. Integrated heats were corrected for the heat of dilution and fitted using an independent model implemented in the NanoAnalyze software (version 3.11.0) to determine the dissociation constant (*K_D_*), binding enthalpy (Δ*H*) and binding stoichiometry (*N*). All measurements were performed at least in duplicate.

## Acknowledgements

The authors gratefully acknowledge support from the European Molecular Biology Laboratory. P.C. was supported by the EIPOD postdoctoral programme co-funded by the European Molecular Biology Laboratory (EMBL) and the Marie Curie Actions Cofund grant MSCACOFUND-FP no. 664726. J.H. gratefully acknowledges the German Research Council (Deutsche Forschungsgemeinschaft, DFG) for support (Grant No.: 267437786, 508497078).

