## Supplementary Figures for "Transient interdomain interactions shape the conformational ensemble governing RNA recognition by the tandem RRMs of Sex-lethal"

**Supplementary Figure 1**

**A**: Spin relaxation data of Sxl-WT (black), Sxl-GS10 (blue), Sxl-GS20 (red) and Sxl-GS25 (green), measured at 600 MHz.

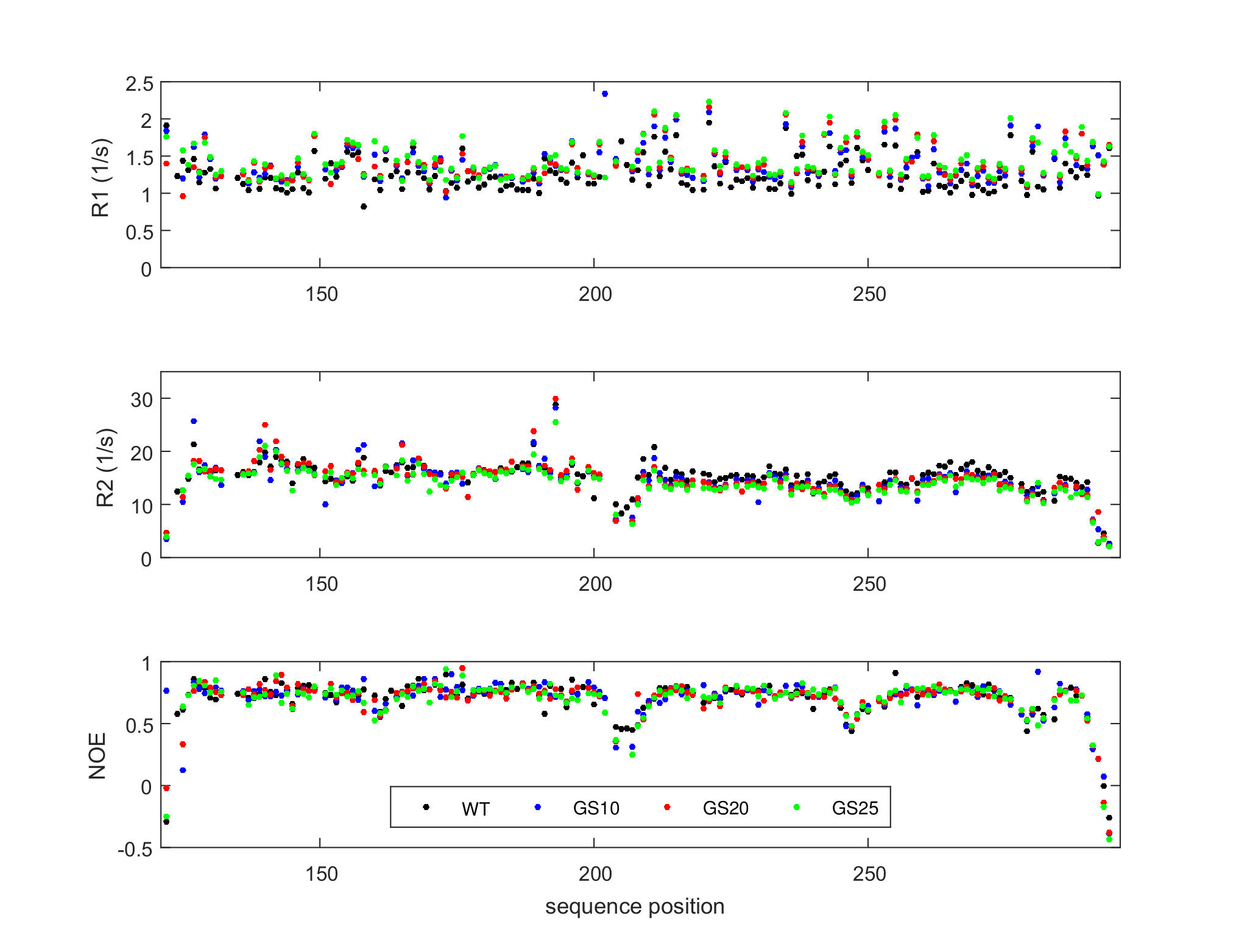

**B**: Spin relaxation data of Sxl-WT (black), Sxl-GS10 (blue), Sxl-GS20 (red) and Sxl-GS25 (green), measured at 1000 MHz.

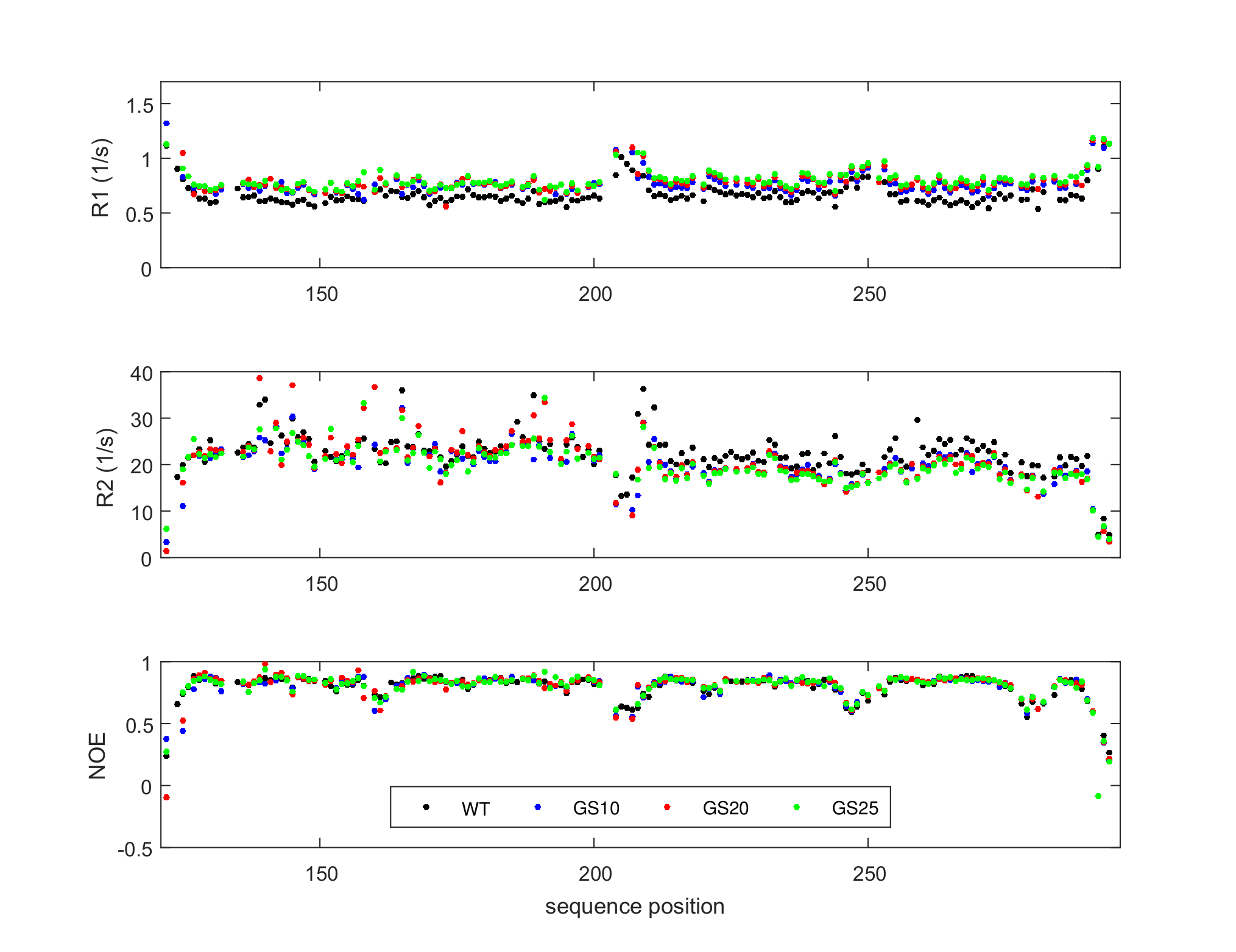

**Supplementary Figure 2**

**A**: Spin relaxation data of Ub-RRM2-GS3 (black), Ub-RRM2-GS5 (blue), Ub-RRM2-GS10 (red) and Ub-RRM2-GS25 (green), measured at 600 MHz.

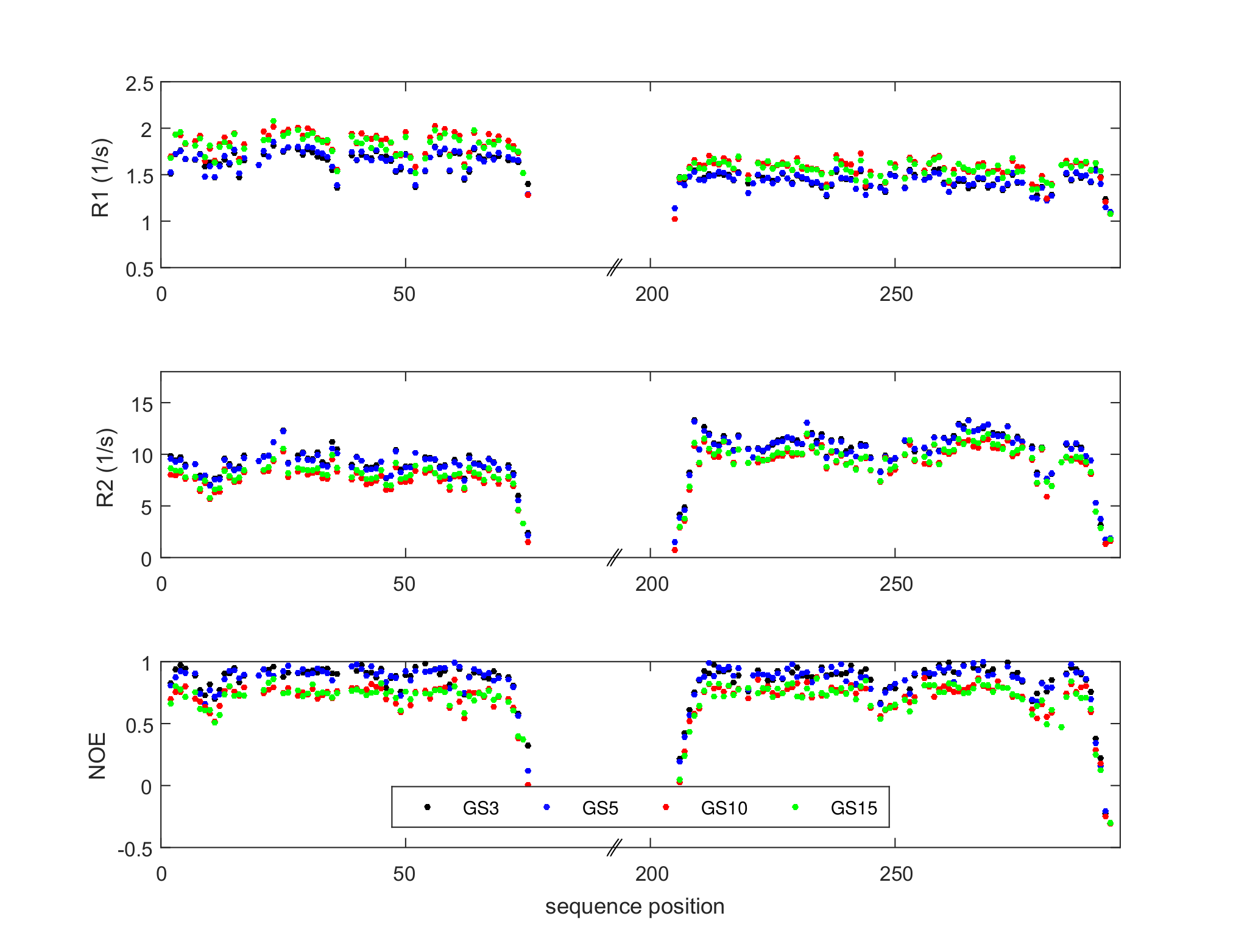

**B**: Spin relaxation data of Ub-RRM2-GS3 (black), Ub-RRM2-GS5 (blue), Ub-RRM2-GS10 (red) and Ub-RRM2-GS25 (green), measured at 800 MHz.

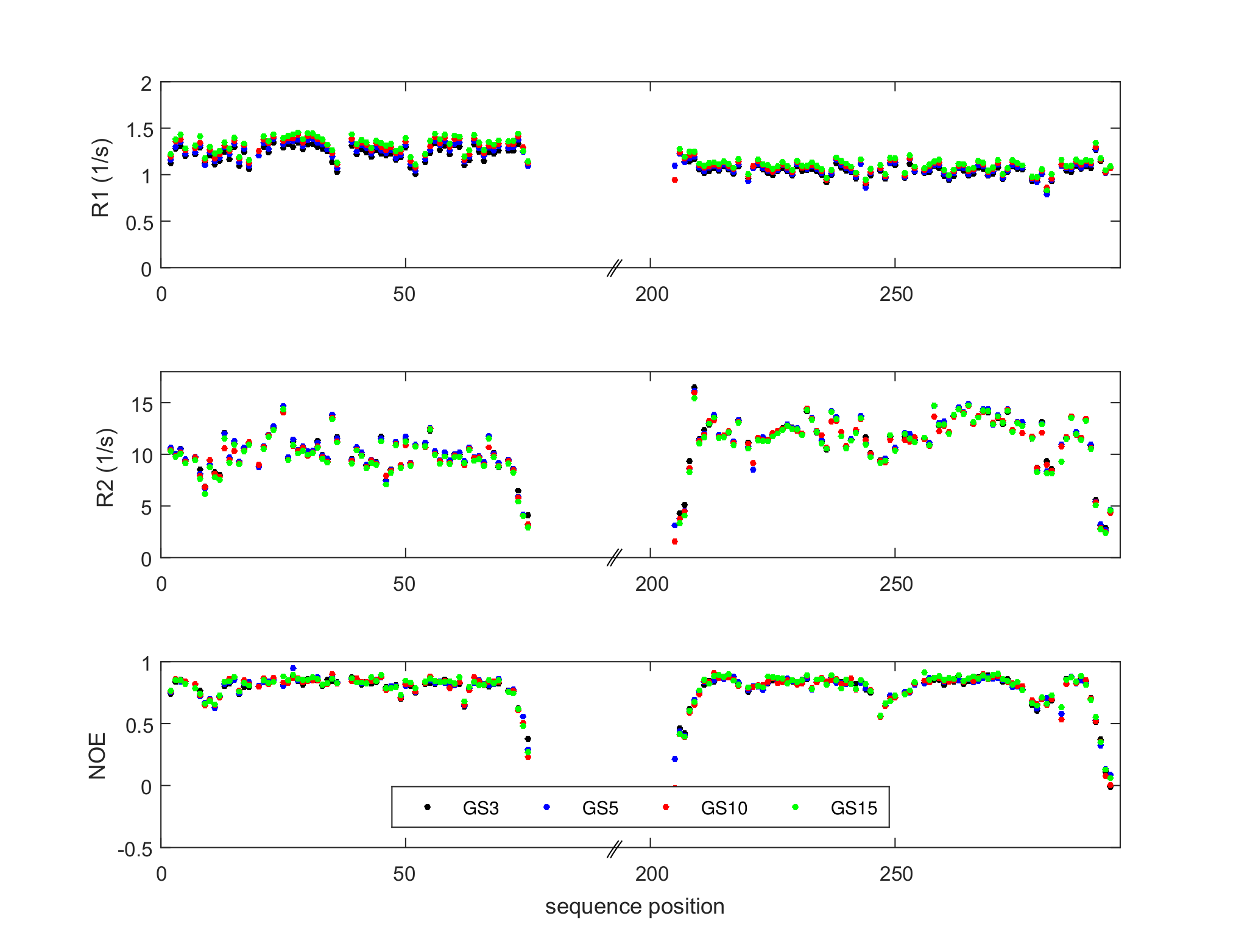

**Supplementary Figure 3**

**A:** ^1^H-^15^N HSQC spectra illustrating linker length-dependent chemical shift changes. Spectra of Sxl-WT, Sxl-GS10, Sxl-GS20, Sxl-GS25, and the isolated RRM1 and RRM2 domains are overlaid as indicated. Colour code is the same as in Figure 4A in the main text (Sxl-WT: black, GS10: purple, GS20: light pink, GS25: magenta, RRM1: light green, RRM2: light blue).

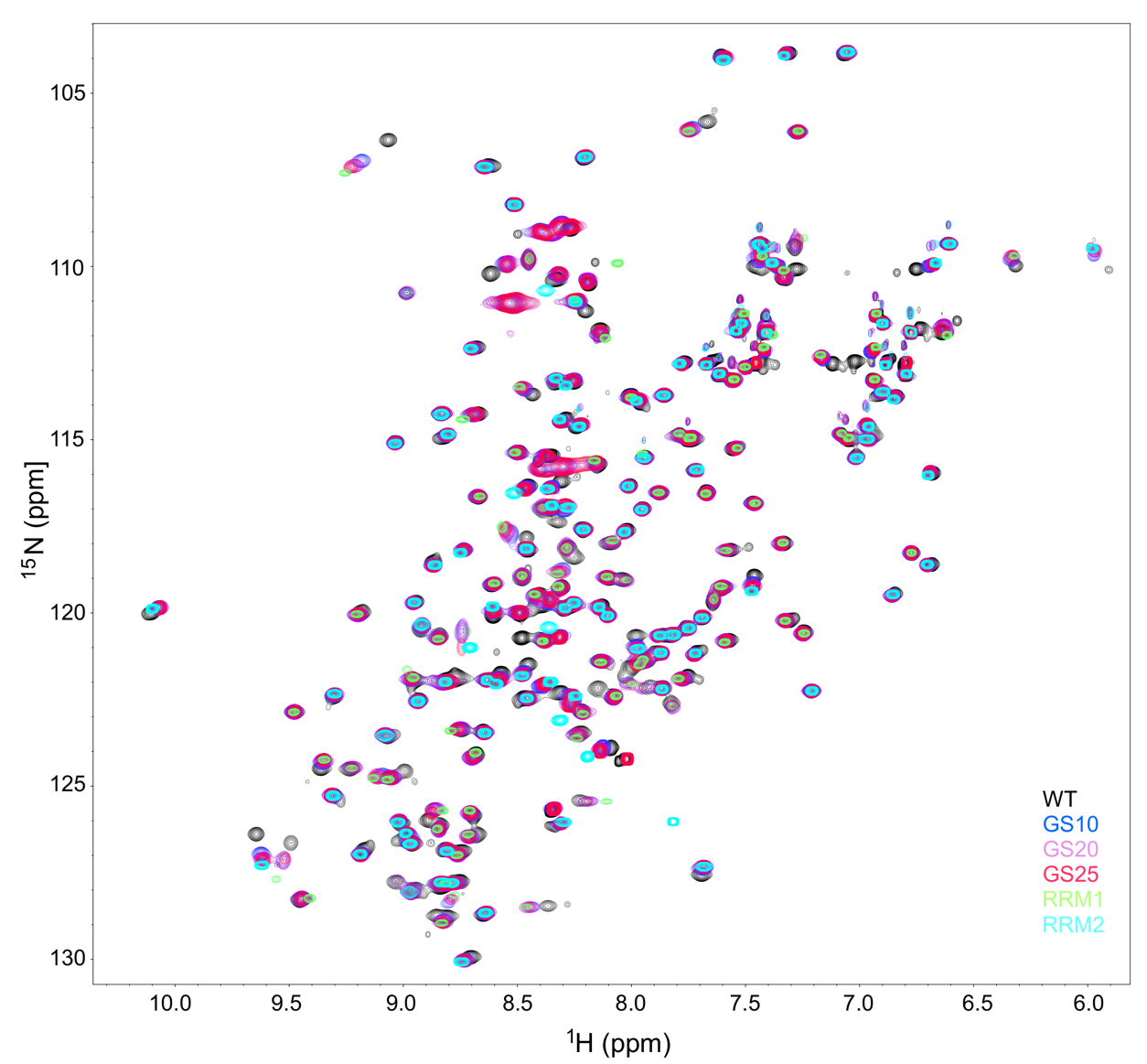

**B:** Amide chemical shift differences plotted as a function of sequence position for each construct and U10-mer RNA binding compared to Sxl-WT. The construct is indicated in the top right corner of each histogram.

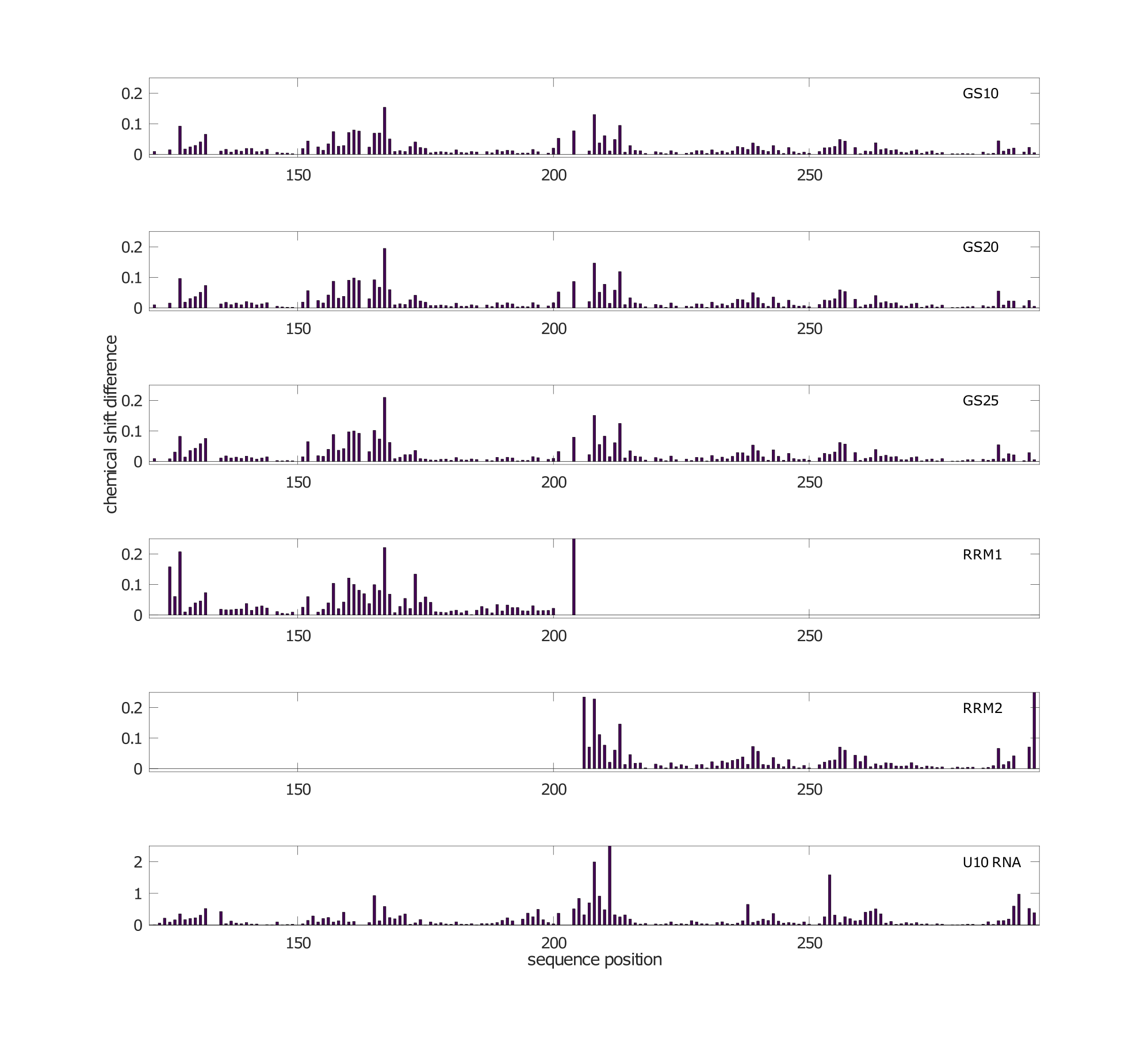

**Supplementar Figure 4**

**A:** Spin relaxation data of Sxl-TYRE mutant (blue) compared with Sxl-WT (black) and Sxl-GS25 (green) measured at 700 MHz.

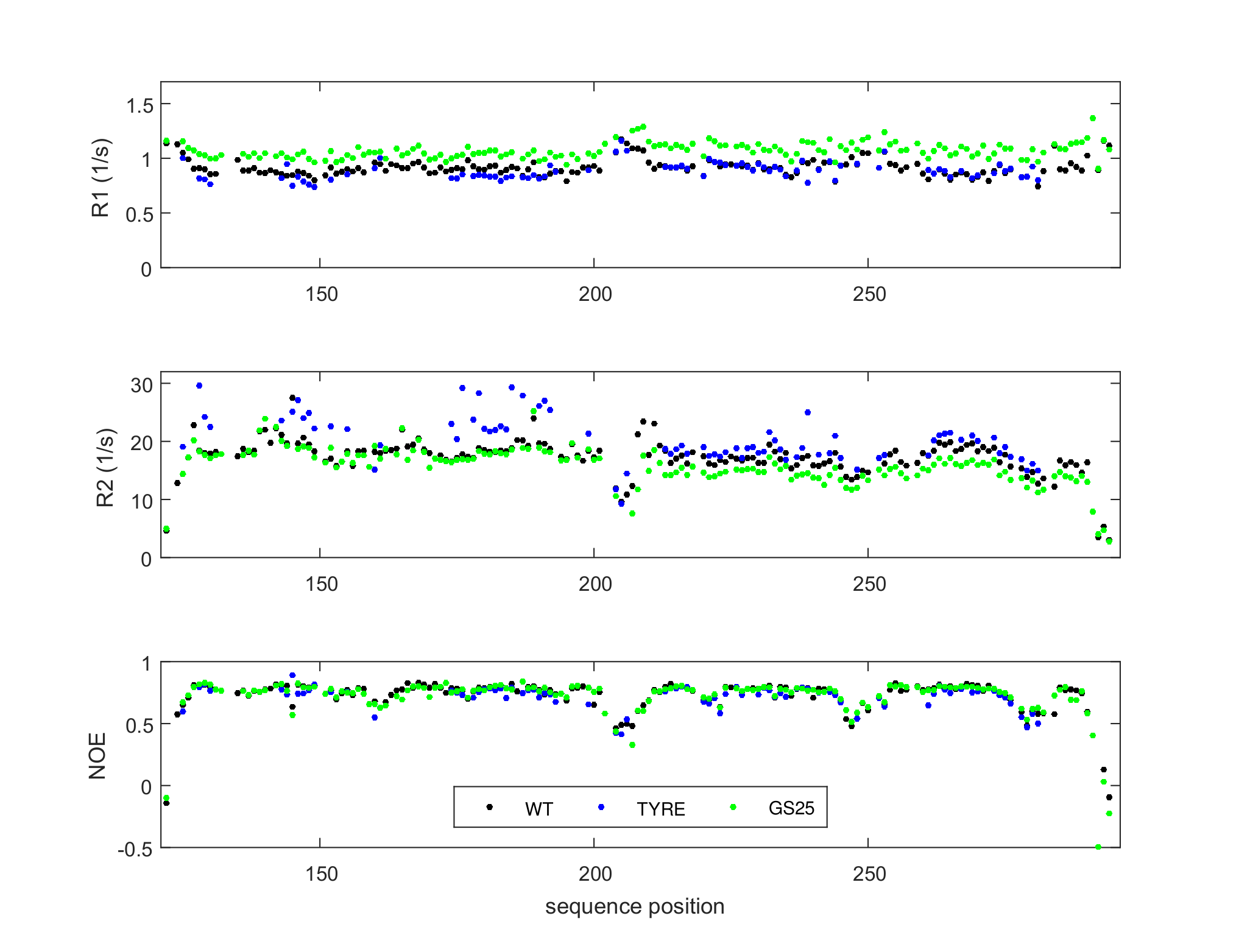

**B:** Full ^1^H-^15^N HSQC spectra of Sxl-WT (black), Sxl-GS25 (red) and the TYRE mutant (blue). The spectra exhibit excellent chemical shift dispersion and comparable peak patterns, demonstrating that all three proteins are properly folded.

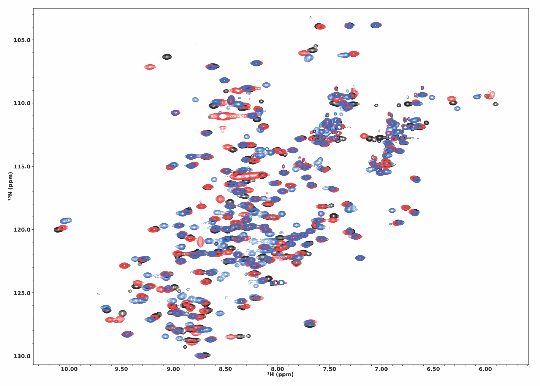

**Supplementary Figure 5**

^1^H-^15^N HSQC spectra of Sxl-WT and Sxl-TYRE mutant titrated with the U5C5 control RNA.

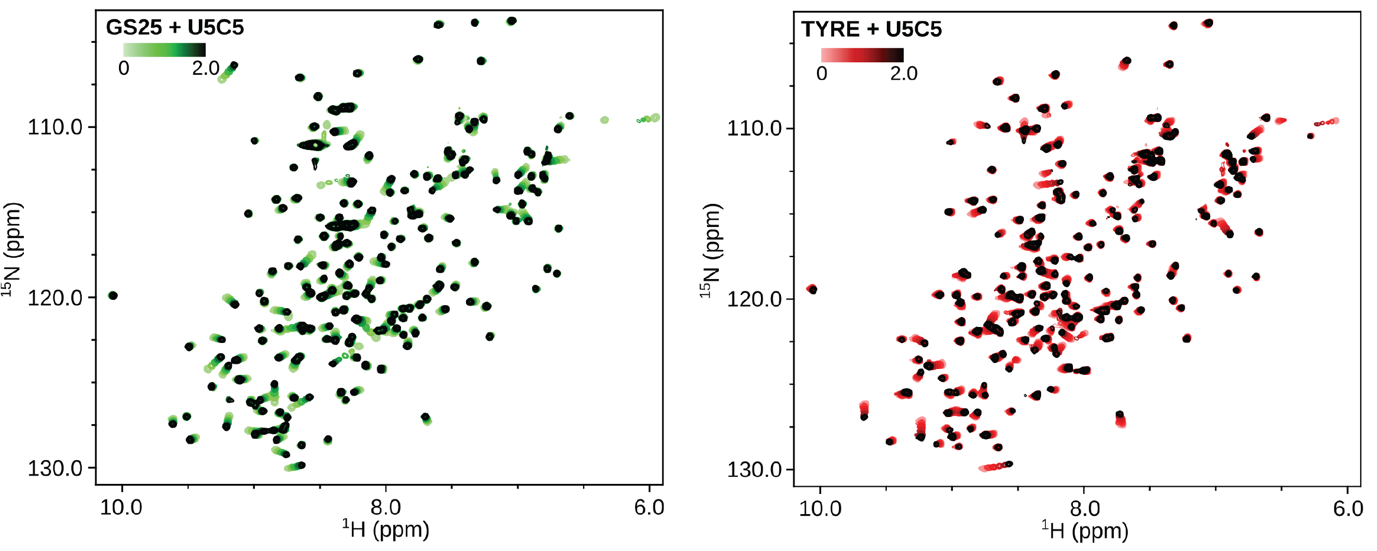
